# Land, carbon and biodiversity data for supply chain impact calculations

**DOI:** 10.1101/2023.11.01.565036

**Authors:** Francis Gassert, Biel Stela, Elena Palao, Mike Harfoot

## Abstract

Monitoring, halting and reversing land conversion is fundamental to meeting international biodiversity and climate targets, and agriculture is the major driver of land conversion. We present an open access set of global data for calculating land use change impacts of agricultural supply chains. These data, originally prepared for the LandGriffon service, include indicators of deforestation, conversion of natural ecosystems, greenhouse gas emissions, and loss of intact or high integrity ecosystems following international standards and guidelines for reporting and target setting in the agriculture, forestry, and land use sector. In order to assign impacts to agricultural production, we prepare data using a spatial adaptation of the statistical Land Use Change (sLUC) accounting approach distributing impact to human activities across the local area using a 50km radius. The results are high resolution global maps of impact per hectare of land occupation. These can then be combined with land footprint data, cropland extent, or productivity maps to calculate land use change related impacts for specific crop volumes sourced from specific regions. Carbon and deforestation results are validated against FAO statistics at the national level.

## Introduction

Human use of land is extensive, it has directly transformed ecosystems across more than 75% of the Earth’s ice-free land area (Ellis 2021). Land conversion, primarily driven by agricultural expansion (Curtis et al. 2018; Seymour and Harris 2019), is one of the foremost drivers of biodiversity loss (IPBES 2019) especially in tropical countries (Harfoot et al. 2021). Global emissions of greenhouse gasses from conversion to agriculture (3.2 GtCO_2_eq, Tubiello et al. 2021) made up 56% of all land use change land use and forestry emissions in 2018 (5.7 GtCO_2_eq, Minx et al. 2021) or 5.5% of global anthropogenic greenhouse gas emissions (58 GtCO_2_eq, Minx et al. 2021). Conversion of natural ecosystems also has major impacts on water cycles and the livelihoods of communities, including indigenous groups, who depend on those ecosystems.

Halting and reversing land conversion and, in particular, deforestation is a fundamental component of international biodiversity and climate targets including those agreed under the Kunming-Montreal Global Biodiversity Framework and Paris Accord. This importance was reflected in the commitments of 141 countries at the UN Climate Conference of Parties (COP26) in Glasgow to halt and reverse forest loss and land degradation by 2030 (UKCOP26 2021). It is also reflected in guidance for companies from the Science Based Targets Network (SBTN, Science Based Targets Network 2023) and the Taskforce for Nature Related Disclosure (TNFD 2023) to report and set targets to halt conversion of natural ecosystems.

Despite this recognition, substantial progress needs to be made. A recent assessment of the 500 companies and financial institutions with the greatest exposure to deforestation risk shows that 40% have not set a single policy on deforestation (Thomson and Fairbairn 2023). And for those that do have a comprehensive commitment, only half are actively monitoring their suppliers or sourcing regions in line with these policies. The lack of readily available datasets on natural ecosystem conversion attributable to agricultural land use contributes to the slow progress towards comprehensive target setting and monitoring.

Here we aim to address this gap by presenting an open access dataset containing preprocessed data for calculating land use change related impacts of agricultural supply chains. We compute complementary indicators that are intended to align with international nature standards and guidelines for reporting, and target setting in the agriculture, forestry, and land use sector. The indicators are designed to allow the attribution of impacts to a land footprint, for example of a site or the land area used to produce a raw material. They were originally developed for use with the Landgriffon service (Harfoot, Palao, and Gassert 2023), which allows a company to ingest its agricultural sourcing data, produce estimates of the resulting environmental impacts and explore scenarios to reduce impacts. The datasets enable organizational reporting, prioritization of actions and resources, and target setting in alignment with Zero Deforestation and land conversion commitments associated with the Accountability Framework Initiative (AFI), TNFD and SBTN.

## Methods

We present a set of global datasets of impacts of commodity production related to human land use following guidance from the Greenhouse Gas Protocol (Greenhouse Gas Protocol 2022), Science Based Targets Initiative (Science Based Targets Initiative 2022) and Science Based Targets Network (Science Based Targets Network 2023). The five land use change related impacts are: deforestation, greenhouse gas emissions from deforestation, net cropland expansion in natural ecosystems, forest integrity or biodiversity intactness of natural ecosystems converted to cropland.

We approach the measurement of conversion of natural ecosystems in two ways, based on the availability of global remotely sensed data on human activities and the natural environment. We map forest loss, excluding areas that are unlikely to be deforestation. We also map cropland expansion into natural ecosystems. Where relevant, we use the SBTN Natural Lands dataset (Mazur et al. 2023) as what we believe to be the best current dataset describing the extent of natural ecosystems.

### Spatial allocation of impacts

The general approach for calculating indicators has two steps. First, we map the spatial extent of natural ecosystem conversion or consequent impacts. Second, we distribute the impacts across all human land using a spatial adaptation of the statistical land use change (sLUC) proportional allocation based on land occupation approach (Greenhouse Gas Protocol 2022).

This *spatial sLUC* approach distributes responsibility for impacts to human land uses over the local area using a kernel radius instead of across the entire country or jurisdiction, such that areas immediately adjacent to land use change receive more responsibility than areas that are far away. This recognizes that commodity-driven land use change occurs outside of existing farms, and that land use adjacent to the forest boundary adds to the land pressure that is driving land conversion.

Specifically, for each pixel, *i*, we compute the total impact, *I*, (e.g. hectares, tons of CO_2_, or index score) within a 50km radius, *r*, and divide it by the total area of non-natural land use, *A*_*nin-natural*_ (Mazur et al. 2023) within the same radius. This gives the ratio of impact per hectare of land occupation, *RI*, in the neighborhood of pixel *i* (Eq. 1, Figure 1). Areas of open water and built areas are considered unavailable to agriculture and are excluded.

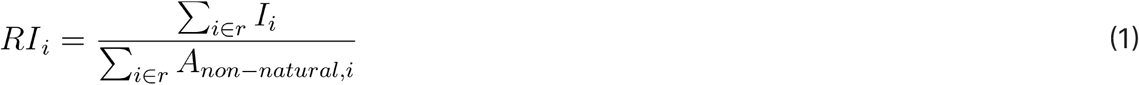

**Figure 1.**
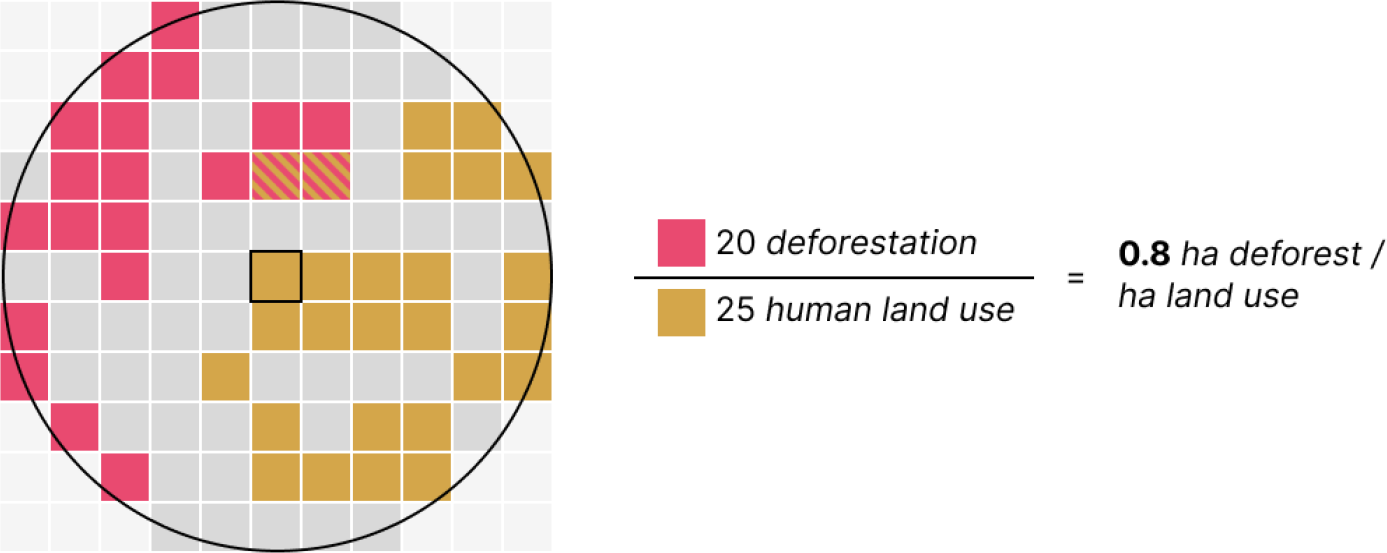
Illustration of sLUC proportional allocation method using a radius around a pixel. Land use produces direct or indirect pressure for land expansion in the surrounding area. For each pixel, we compute the total area of deforestation within a given radius and divide by the total area of non-natural land use to allocate responsibility for deforestation. The result is that land use in close proximity to land expansion boundary receives a greater responsibility for deforestation.

If this ratio of land conversion to land occupation exceeds 1.0 (i.e. there is more deforestation than human land use) it is capped at 1, such that a single hectare of human land use cannot be responsible for more than one hectare of land conversion and associated impacts.

Deforestation related indicators are annualized for the previous 20 year period (2002-2022) to align with SBTi guidance on carbon. Cropland expansion indicators are annualized over the two year period following 2020 to align with SBTN guidance on conversion of natural ecosystems.

All computation is performed in Google Earth Engine in the WGS84 projection. Raster math is performed at a 100m (0.000898 degree) resolution. Kernel convolutions are performed at a 1km (0.00898 degree) resolution to reduce computational cost.

### Indicator methodologies

The following sections detail the methodologies used for calculation of the impact, *I*, for each of the five impact indicators.

#### Deforestation

The deforestation indicator estimates the extent of deforestation that can be attributed to a spatial footprint, such as that required for production of a raw material. It measures the amount of deforestation that occurs nearby to the areas where raw materials are produced within a supply shed.

Deforestation is the conversion of forested areas to non-forest land use such as arable land, urban use, logged area or wasteland (FAO 2023). We estimate the extent of deforestation using data from Global Forest Watch by looking first at all areas of tree cover loss over the past 20 years (Hansen et al. 2013), and then exclude the following areas where tree cover loss is unlikely to be a change in land use:

1. Loss of non-natural tree cover outside of areas classified as intact forests in 2000. Non-natural tree cover is identified from (Mazur et al. 2023) which uses the Spatial Database on Planted Trees (Richter et al. in review) to classify tree crops (c.2010-2020) as non-natural. Intact forests are identified using Intact Forest Landscapes (Potapov et al. 2017) and Primary Humid Tropical Forests (Turubanova et al. 2018). This is intended to exclude harvesting within plantation woodlots and changes in tree cropping patterns.
2. Forest disturbance (Potapov et al. 2022) outside of areas classified as intact forests in 2000. Forest disturbance identifies areas with trees of >5m in height in both 2000 and 2020, but experienced significant disturbance in the intervening period. This is intended to exclude long-standing land use patterns of rotational logging and shifting agriculture, as well as natural forest disturbances (wildfire, blowdown) in secondary forests.
3. Tree loss that corresponds with burned areas outside of tropical and subtropical biomes (Tyukavina et al. 2022). This is intended to exclude tree cover loss due to wildfire in biomes where fire is a frequent natural occurrence.

#### Greenhouse Gas Emissions from Deforestation

Deforestation carbon loss is the amount of carbon emissions that can be associated with a deforestation event. We estimate forest carbon loss due to deforestation by combining deforestation extent with maps of vulnerable carbon (Noon et al. 2021) in those same areas. Vulnerable carbon, the amount of biomass and soil carbon that would be lost in a land use change event, is reported globally for 2010 at a 300m resolution.

For areas that were deforested between 2000 and 2010, we estimate year-2000 vulnerable carbon by filling gaps in forest carbon in the 2010 data. We select pixels that are classified as forests in 2010 in the ESA CCI Land Cover time series v2.0.7 dataset (ESA 2017), which was used as an input to generate the vulnerable carbon data, and compute mean vulnerable carbon for forests in the local area using a 10km kernel. We assign the mean vulnerable carbon values for forest to pixels that are classified as forests in the year 2000 by Hansen et al. (2013), but not classified as forests in 2010 by ESA. This ensures that all pixels that may be identified as deforested have a carbon value that is reflective of the local area.

#### Net cropland expansion in natural ecosystems

Due to limitations in remotely sensed data on sparse human land uses such as logging and pasturing, we use a conservative approach evaluating only the conversion of natural ecosystems directly to cropland, using the SBTN Natural Lands dataset (Mazur et al. 2023).

We assess cropland expansion using the Esri LC 10m cropland class (Karra et al. 2021). We consider any area that is classified as cropland in one any of the previous three years to be cropland to account for temporal variations (i.e. fallowing) and minimize inaccuracies. E.g. such that any cropland between 2017 and 2020 is considered cropland for the year 2020.

To identify natural ecosystem conversion to cropland, we first identify the total land area that is considered to be cropland in 2022 but not in 2020, filtered to regions that are classified as natural in SBTN Natural Lands dataset. We also identify cropland reduction as areas that are cropland in 2020 but not in 2022. Finally, for each pixel, we compute the total area of crop expansion in natural ecosystems within 50km, and the total area of cropland reduction in natural ecosystems within 50km. We compute net cropland expansion in natural ecosystems as expansion minus reduction, excluding areas with more reduction than expansion.

#### Loss of forest integrity and biodiversity intactness to cropland expansion

There are many facets of biodiversity that vary in importance for different stakeholders. For this reason several biodiversity risk indicators are proposed reflecting these different aspects. To align with previous analysis and with guidance from the SBTN, two categories of indicator are often proposed, one focussed on species and the other on ecosystems. Here we specifically use an ecosystem focussed indicator.

We compute two indicators of that attempt to capture the potential loss of biodiversity due to land use change. Each indicator evaluates the level of naturalness or intactness of the ecosystems that were lost to cropland expansion.

We use estimates of forested ecosystem intactness using the Forest Landscape Integrity Index (Grantham et al. 2020) and from the Biodiversity Intactness dataset (Gassert, Mazzarello, and Hyde 2022). The Forest Landscape Integrity Index (FLII) expresses the degree of intactness of forest ecosystems based on the spatial structure of the forest habitat and on the degree of human modification to the ecosystem. The Biodiversity Intactness index (BI) is an indicator of loss of species richness and change in species composition due to human pressures. Notably, FLII takes a composite indicator approach to estimate integrity, is limited to forest ecosystems, and takes into account the level of fragmentation or connectedness of the ecosystem. BI on the other hand attempts to model the relationship between human pressures and the loss of biodiversity using mixed effects regression on site level data from the PREDICTS dataset (Hudson et al. 2017).

For both indicators, we compute the sum of the index for natural ecosystems that were converted to croplands within the 50km radius. As both of these datasets report results as unitless scores, they are primarily useful for prioritization or comparison.

### Data record

Datasets are provided under a Creative Commons Attribution 4.0 International License at https://doi.org/10.5281/zenodo.10048050. This repository includes gridded data products for each of five indicators:

- Indicators:
  - Deforestation
  - Greenhouse gas emissions from deforestation
  - Net cropland expansion in natural lands
  - Loss of forest integrity to cropland expansion
  - Loss of biodiversity intactness to cropland expansion
- Layers:
  - Impact (100m resolution)
  - Impact per hectare of land occupation (1km resolution)

Scripts to produce these datasets and validation analysis are available on GitHub at https://github.com/vizzuality/lg-landscape-indicators-processing.

## Results and interpretation

Results are provided as gridded datasets of impact per hectare of human land occupation, such that users can multiply this data by the land footprint of commodity production or other human activities to calculate the impact of sourcing specific raw materials from that location (Figure 2). Figure 3 shows the impact indicators clipped to all non-natural areas.

**Figure 2.**
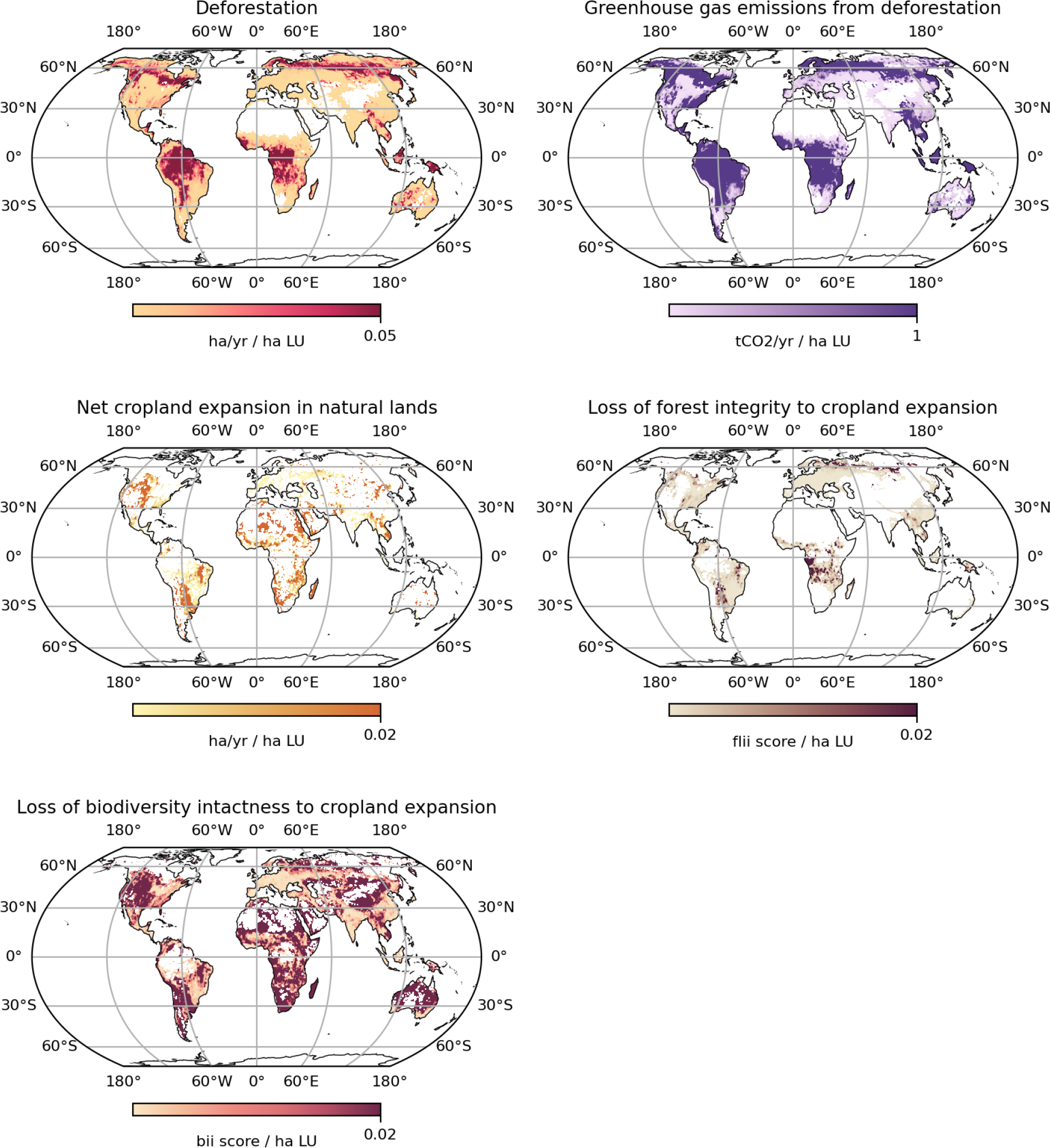
Maps of impact per hectare of land occupation (LU). Higher values show that land use in those regions would receive higher impact scores. These maps are intended to be combined with spatial data on the land footprint of agricultural or forestry production to calculate the impacts associated with supply chain sourcing. They are not constrained to areas of current human land use, for example a 1ha of land use in the middle of remote forest would be assigned a maximum impact of 1ha deforestation amortized over 20yrs.

**Figure 3.**
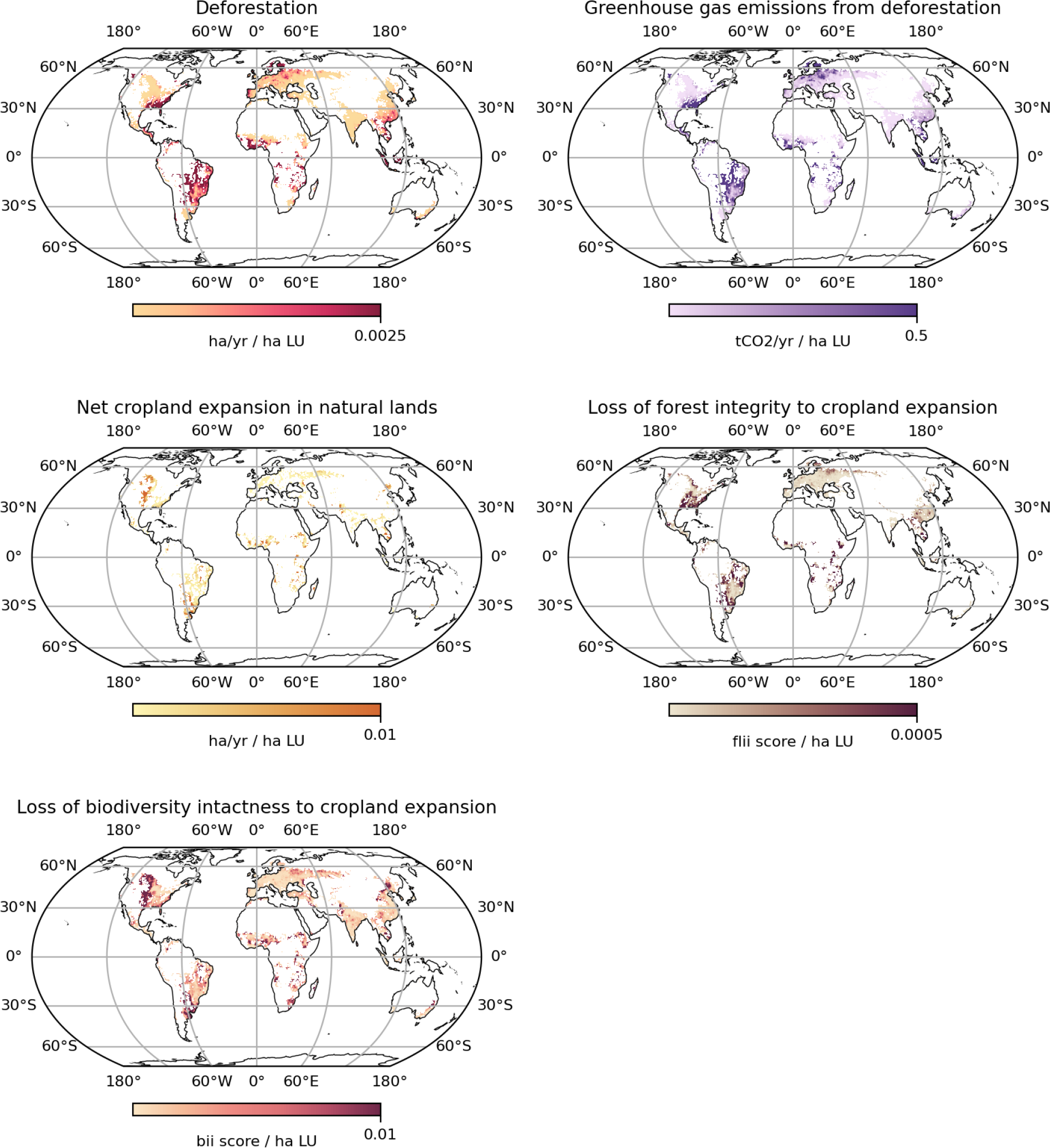
Maps of impact per hectare of land occupation (LU) masked to areas of non-natural (human) land use. These maps represent the spatial distribution of impacts that can be attributed to land occupation following the spatial sLUC approach.

Methods for combining these results with spatially explicitly production data and land footprint data to calculate the impacts of raw materials sourced from for local, regional, or national supply-sheds are documented in the LandGriffon methodology (Harfoot, Palao, and Gassert 2023). For example, to calculate the deforestation footprint of sourcing 100 tons of soy from Argentina, a user could first calculate the average hectares of deforestation per hectare of land occupation for all soy producing areas in Argentina, and then multiply that value by the land footprint (reciprocal of crop yield) of 100 tons of soy production.

### Validation

We are not aware of spatially explicit estimates of deforestation areas or greenhouse gas emissions associated with land conversion with which we could evaluate our sLUC data layers. We therefore chose to validate deforestation area and greenhouse gas emissions associated with deforestation by comparing our results with statistics from the UN Food and Agricultural Organization (FAO) over the same 20-year period. We compare results for total deforestation area and carbon by country, as well as by the proportion that can be attributed to cropland expansion.

For our datasets, we attribute impacts to cropland using the statistical sLUC method for all areas identified as cropland by Potapov et al. (2021). For FAO statistics we use the ratio of nationally reported cropland area to that of all agricultural land to allocate impacts to cropland. We are not aware of extant ground truth data for the conversion of natural ecosystems to cropland, and did not attempt to collect our own data for validation. Likewise, we do not attempt to validate the biodiversity impact indicators.

We report validation results for the 189 for countries for which the FAO reports forest conversion statistics and for the subset of 118 tropical countries (Table 1, Figure 4). For tropical countries, our results are comparable to FAO statistics despite differences in data and methods. Pearson’s coefficient of determination (r^2^) for total deforestation area was 0.95 and for related GHG emissions of 0.93. Results are substantially less aligned when including temperate regions, with our data showing higher deforestation area and related emissions than FAO statistics (r^2^ < 0.3). Results for the portion of deforestation area and related GHG emissions attributable to cropland are less aligned than for country totals.

**Table 1.**
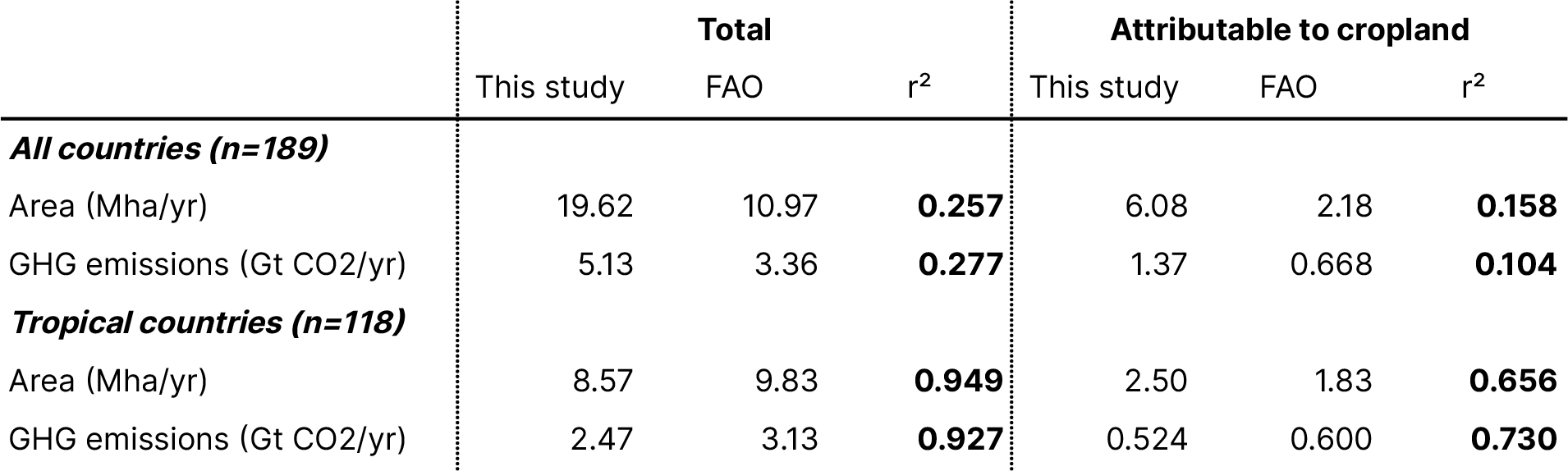
Validation results comparing total area of forest conversion and associated greenhouse gas emissions from this study vs from FAO statistics for 189 countries with reported forest conversion data in FAO.

|  | Total |  |  | Attributable to cropland |  |  |
| --- | --- | --- | --- | --- | --- | --- |
|  | This study | FAO | r<sup>2</sup> | This study | FAO | r<sup>2</sup> |
| **All countries (n=189)** |  |  |  |  |  |  |
| Area (Mha/yr) | 19.62 | 10.97 | **0.257** | 6.08 | 2.18 | **0.158** |
| GHG emissions (Gt CO2/yr) | 5.13 | 3.36 | **0.277** | 1.37 | 0.668 | **0.104** |
| **Tropical countries (n=118)** |  |  |  |  |  |  |
| Area (Mha/yr) | 8.57 | 9.83 | **0.949** | 2.50 | 1.83 | **0.656** |
| GHG emissions (Gt CO2/yr) | 2.47 | 3.13 | **0.927** | 0.524 | 0.600 | **0.730** |

**Figure 4:**
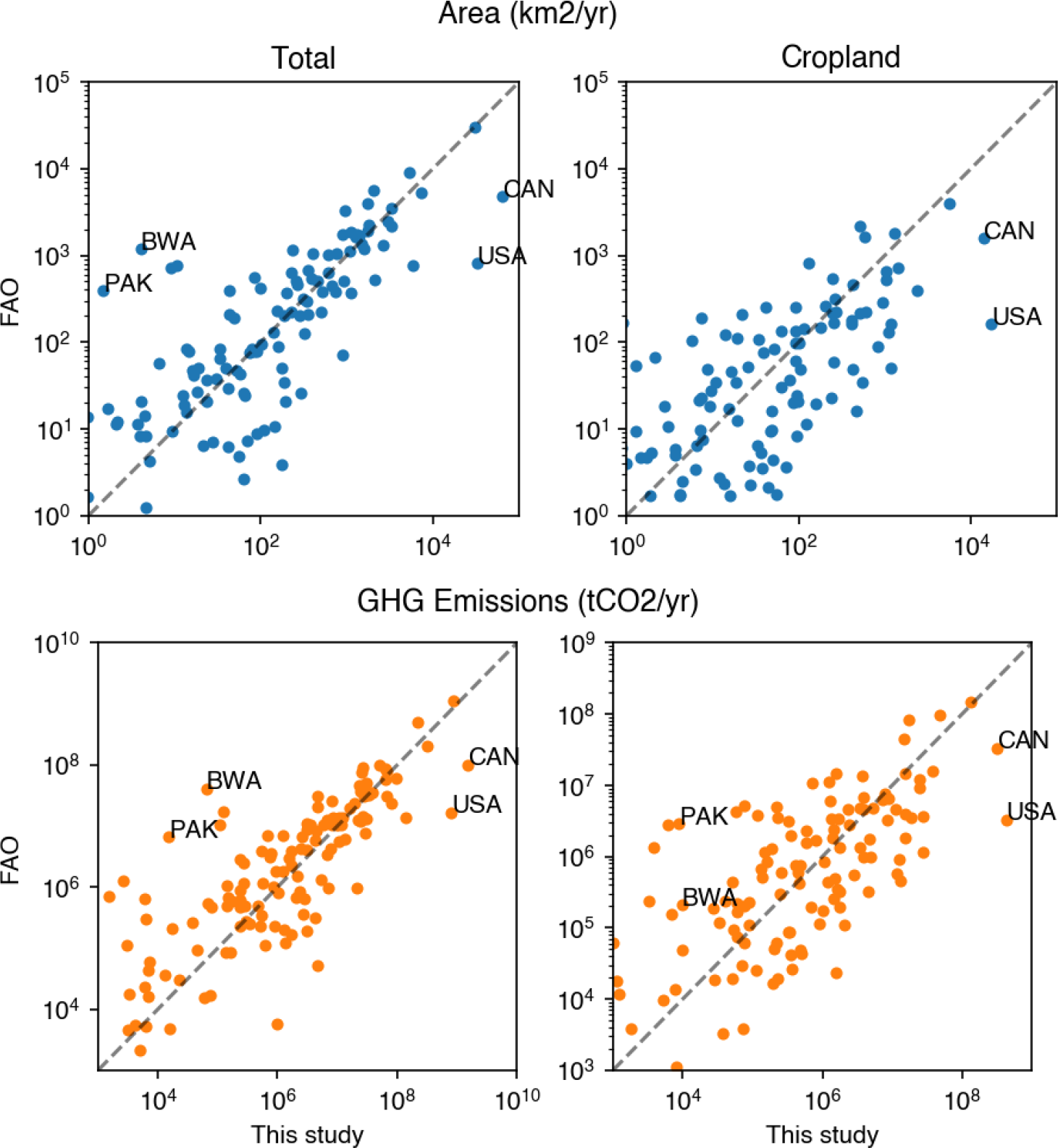
Comparison of forest conversion area and related greenhouse gas emissions results from this study against FAO statistics for 189 countries reporting forest conversion statistics to FAO. Notable outlier countries are labeled.

## Discussion

The datasets made available in this study make spatially explicit attribution of impacts to land use substantially more accessible for organizations with land footprints. The democratization of this impact calculation is of great value for agricultural supply chain sustainability management. The results also open the possibility for novel analyses of where land conversion, especially to cropland, is driving most natural ecosystem conversion, and therefore where activities to reduce expansion could have the most benefit.

### Interpretation

Our results for deforestation and associated GHG emissions show reasonable alignment with official FAO statistics, and substantially better agreement than raw forest loss estimates. However, we acknowledge that there are notable outliers in comparison with official statistics. In particular, we substantially overestimate deforestation in the U.S. and Canada, these two countries together account for most of the difference in total area of deforestation and related emissions relative to official statistics. This is likely due to the practice of rotational logging with natural regrowth, as we will classify areas of recent rotational logging of secondary forests where the area had not regrown as of 2020 as deforestation when they may not be a true land use change. Areas with plantation woodlots will be less affected by this issue as we seek to specifically exclude them from our deforestation statistics.

Conversely, we underestimate deforestation in countries such as Botswana and Pakistan. This is likely due to the difficulty detecting sparse and dry forests in the remotely sensed forest cover change data.

Results for the portion of deforestation attributed to cropland show a lower degree of alignment. However, this should be expected as we account for the spatial distribution of cropland within countries, when this is not possible when calculating from national level statistics.

### Usage notes

As a result of the above interpretation, users of our deforestation data should bear in mind that it is likely to overestimate forest conversion in areas of long-standing rotational forestry, and underestimate deforestation in areas of sparse and dry forest. Results for the conversion of natural ecosystems into cropland are unlikely to have the same issues, but may suffer from other unknown biases.

These data are intended to be used to estimate impacts related to the production of commodities in supply chains. They provide a free and open-access, globally consistent and earth-observation based source of data, designed to be updated through time as new data becomes available. This time dynamic aspect is particularly important to track impacts and prioritize materials, suppliers, and regions where additional data collection, due diligence, or supplier engagement may be warranted.

Importantly, these data are not intended to replace official statistics and should not be used to estimate national greenhouse gas footprints. We do incorporate national level data, such as higher accuracy land use change maps or data on forest concessions which are necessary to understand the true scope of land conversion. Moreover, these data do not capture greenhouse gas emissions and biodiversity impacts associated with non-deforestation events such as wildfire, shifting agriculture, and wood harvesting.

### Conclusion

Spatial sLUC approach has the advantage of having a closer spatial relationship with actual sourcing regions relative to country level statistics, while not requiring the identification of farm boundaries or specific production locations. It provides consistent spatial assumptions regardless of the area of analysis, which enables users to calculate impacts for supply sheds as small as individual farms or as large as countries or regions and get consistent and comparable results. In addition, while we do not attempt to map the expansion of pasture lands, the spatial sLUC approach captures the indirect land use impacts of cropland expansion may cause by displacing other land users.

If we are to meet our pressing international biodiversity and climate targets, we need organizations to act now to reduce land conversion. These datasets provide open, and actionable data that allow organizations to monitor where they are individually or collectively contributing to land conversion and take action to reduce these impacts.

Ultimately, society needs more such data to permeate decision-making and drive rapid action to halt the expansion of human land use and its resultant impacts on biodiversity and the Earth system.

